# Structural-Kinetic-Thermodynamic Relationships Identified from Physics-based Molecular Simulation Models

**DOI:** 10.1101/183053

**Authors:** Joseph F. Rudzinski, Tristan Bereau

## Abstract

Coarse-grained molecular simulation models have provided immense, often general, insight into the complex behavior of condensed-phase systems, but suffer from a lost connection to the true dynamical properties of the underlying system. In general, the physics that is built into a model shapes the free-energy landscape, restricting the attainable static and kinetic properties. In this work, we perform a detailed investigation into the property interrelationships resulting from these restrictions, for a representative system of the helix-coil transition. Inspired by high-throughput studies, we systematically vary force-field parameters and monitor their structural, kinetic, and thermodynamic properties. The focus of our investigation is a simple coarse-grained model, which accurately represents the underlying structural ensemble, i.e., effectively avoids sterically-forbidden configurations. As a result of this built-in physics, we observe a rather large restriction in the topology of the networks characterizing the simulation kinetics. When screening across force-field parameters, we find that structurally-accurate models also best reproduce the kinetics, suggesting *structural-kinetic relationships* for these models. Additionally, an investigation into thermodynamic properties reveals a link between the cooperativity of the transition and the network topology *at a single reference temperature*.

## I. INTRODUCTION

Coarse-grained (CG) molecular simulation models not only allow accelerated investigations of condensed-phase systems, but provide a means to understand the essential driving forces for particular structures and processes. More specficially, CG models are an interesting test bed for gaining insight into the relationship between particular force field components and the resulting features of the free-energy landscape. The beneficial speed-up of CG models, attained through a combination of reduced molecular friction and softer interaction potentials, comes at the cost of obscuring the connection to the true dynamical properties of the underlying system. Unlike many polymer systems, where a homogeneous dynamical rescaling factor is capable of recovering the correct dynamics of the underlying system,^1,2^ the rescaling of CG dynamical processes may generally be a complex function of the system’s configuration. This lost connection to the true system dynamics represents a severe limitation for CG models, which may not only prevent quantitative prediction of kinetic properties, but may also lead to qualitatively misleading or incorrect interpretations generated from CG simulations. For example, Habibi *et al*.^3^ recently demonstrated that three different CG models provide disparate descriptions of the forced unfolding process of a 110 residue peptide, despite the capability of all models to fold the peptide to the proper native structure. Furthermore, the fidelity of the folding process, with respect to an AA reference simulation, did not correlate with the complexity of the model. Perhaps more troubling, Rudzinski *et al*.^4^ recently demonstrated that, even for a tripeptide of alanines, a transferable physics-based CG model qualitatively misrepresents the hierarchy of local dynamical processes along the peptide backbone, i.e., transitions between metastable states on the Ramachandran plot.

Although it is possible in principle to rescue the dynamics of a CG model via a generalized Langevin formalism,^5^ this approach offers a daunting computational and conceptual challenge for complex biological molecules that give rise to hierarchical dynamics, i.e., kinetic processes coupled over various timescales. As an alternative, it may be possible to rescue the kinetic properties generated from a CG simulation via an a posteriori reweighting of simulation data in order to reproduce a set of provided reference observables, such as experimental measurements.^4,6^ However, the extent to which experimental data can correct for deficiencies in simulation models through a reweighting is obviously limited. Thus, a detailed understanding of the link between given target observables, structural properties for instance, and the accurate reproduction of relative kinetic quantities is required. Moreover, because there are established methods for systematically addressing the structural accuracy of CG models,^7^ identification of robust relationships between structural and kinetic properties provides a practical route for improving the dynamical accuracy of CG models for biomolecular systems.

Our previous studies^4,6^ revealed that if there is relevant physics missing from a CG model, for example, conformational states which are sterically forbidden in the true ensemble but sampled by the CG model, the kinetic properties may demonstrate drastic errors, despite relatively high accuracy in the description of structural properties. In the present study, we investigate this concept in reverse. That is, assuming a model incorporates the essential physics, does reproducing *enough* structural detail lead to accurate reproduction of kinetic properties? More specifically, we investigate relationships between structural and kinetic properties in the helix-coil transition networks generated by various simulation models. As a fundamental process in protein folding, investigation of *α*-helical secondary-structure formation represents a logical step in understanding the kinetic properties generated by CG protein models. Remarkably, we find that the physics of the model enforces implicit constraints on the free-energy landscape, which results in simple connections between structural and kinetic properties—accurate structural features guarantee consistent kinetics. These structural-kinetic relationships ease the interpretation of the CG helix-coil kinetics, justifying the use of a homogeneous speed-up factor. Our approach extends upon previous works that have performed systematic scans of model parameters for relatively simple CG models to obtain insight into the driving forces for emergent condensed-phase properties.^8–10^ Conceptually, this strategy mirrors recent efforts at uncovering structure-property relationships by systematic high-throughput studies,^11–13^ but focuses on probing property interrelationships by simultaneously monitoring various emergent features of the model. As a primary tool for investigation, we employ a native-biased model, whose parameters may be easily tuned to elucidate the connection between specific interactions, e.g., hydrophobic attraction between side chains, and emergent properties of the resulting kinetic network. However, in order to sample a physically-realistic ensemble of structures (likely a necessary condition for the consistent reproduction of kinetic properties), this model retains a near-atomistic description of backbone steric interactions. To assess the generality of the results, we additionally consider a transferable, physics-based CG peptide model, which also retains near-atomistic backbone resolution but employs phenomenological interactions in order to reproduce the balance of *α*/*β* structural propensities in a number of distinct peptide systems.^14,15^

As a model system, we examine an uncapped heptapeptide of alanines in the unfolded regime, where a disordered ensemble of pathways between the helix and coil states gives rise to a complex hierarchy of relevant kinetic processes. We demonstrate that matching particular structural properties of the ensemble fixes the ratio of certain timescales, given the implicit constraints provided by the underlying physics of the model. By characterizing the temperature dependence of the various CG models, we further validate these structural-kinetic relationships while also demonstrating that constraining simple structural properties at a single temperature is not enough to dictate the cooperativity of the resulting transition. However, an investigation of the transition networks characterizing the simulation kinetics reveals a connection between the cooperativity of the model and the network topology at a single reference temperature.

## II. METHODS

### A. Coarse-grained (CG) models

#### 1. Hybrid Gō (Hy-Gō)

To investigate the relationship between structural and kinetic properties generated from CG simulation models, we wanted a simple model (i.e., with few, physically-motivated parameters) which samples all the relevant conformational states. For this purpose, we employ a Gō-type model,^16^ which defines attractive interactions based on the location of atoms in the native structure. In particular, we use a flavored-Gō model with three interaction types:^17–19^ (*i*) a native contact (nc) attraction, *U*_nc_, employed between pairs of *C*_*α*_ atoms which lie within a certain distance in the native structure, i.e., the *α*-helix, of the peptide. (*ii*) a desolvation barrier (db) interaction, *U*_db_, also employed between native contacts, and (*iii*) a hydrophobic (hp) attraction, *U*_hp_, employed between all pairs of C*α* atoms of the amino acid side chains. *U*_nc_ and *U*_hp_ are necessary for sampling the correct ensemble of conformations (i.e., helix, coil, and swollen structures), while *U*_db_ assists in providing cooperativity in the resulting transitions. We employed the same functional forms as in many previous studies,^19^ with a tunable prefactor for each interaction as described below.

These three Gō-type interactions result in models which *roughly* sample the correct conformational ensemble for short peptides. However, because we are interested in characterizing kinetic properties, it is important that we reproduce the underlying conformational ensemble more precisely. More specifically, we wanted to ensure that the model (*i*) does not sample conformations that are sterically forbidden in the all-atom (AA) model and (*ii*) samples the relevant regions of the Ramachandran plot, while retaining barriers between metastable states. Thus, in addition to the simple interactions described above, the model also partially employs a standard AA force field, AMBER99sb,^20^ to model both the steric interactions between all non-hydrogen atoms and also the specific local conformational preferences along the chain. More specifically, the bond, angle, dihedral, and 1–4 interactions of the AA force-field are employed without adjustment. To incorporate generic steric effects, without including specific attractive interactions, we constructed Weeks-Chandler-Andersen potentials^21^ (i.e., purely repulsive potentials) directly from the Lennard-Jones parameters of each pair of non-hydrogen atom types in the AA model. For simplicity of implementation, we then fit each of these potentials to an *r*^‒12^ functional form. The van der Waals attractions and all electrostatic interactions in the AA force field were not included and water molecules were not explicitly represented.

Although the local sterics of the backbone are modeled with near-atomistic resolution, the Hy-Gō model substitutes the full atomistic description of the peptide with a highly coarse-grained representation by modeling the complex combination of dispersion and electrostatic peptide-peptide and peptide-solvent interactions with a limited set of simple interactions between C_*α*_ and C*β* atoms. Note that this is quite distinct from all-atom Gō^22^ models, which employ atomically-detailed energetic parameters based on the positions of each atom in the native structure. The total interaction potential for the present model may be written: *U*_tot_ = *ϵ*_nc_*U*_nc_ + *ϵ*_db_*U*_db_ + *ϵ*_hp_*U*_hp_ + *ϵ*_bb_ *U*_bb_, where the backbone (bb) interaction includes both the intramolecular and steric interactions determined from the AA force field. The first three coefficients represent the only free parameters of the model, while *ϵ*_bb_ = 1. The relative impact of of the backbone interactions may then be characterized by *E*_tot_ ≡ *ϵ*_nc_ + *ϵ*_db_ + *ϵ*_hp_.

#### 2. PLUM

As an alternative CG model, we considered the PLUM model, which also describes the protein backbone with near-atomistic resolution, while representing each amino acid side chain with a single CG site, within an implicit water environment.^14^ In PLUM, the parametrization of local interactions (e.g., sterics) aimed at a qualitative description of Ramachandran maps, while longer-range interactions—hydrogen bond and hydrophobic— aimed at reproducing the folding of a three-helix bundle, without explicit bias toward the native structure.^14^ The model is transferable in that it aims at describing the essential features of a variety of amino-acid sequences, rather than an accurate reproduction of any specific one. After parametrization, it was demonstrated that the PLUM model folds several helical peptides,^14,23–26^ stabilizes *β*-sheet structures,^14,15,27–29^ and is useful for probing the conformational variability of intrinsically disordered proteins.^30^ We also considered two minor reparametrizations of the PLUM model: 1. The side chain van der Waals radius is decreased to 90 % of its original value,^6^ and 2. The hydrogen-bonding interaction strength is decreased to 94.5 % of its original value.^30^

### B. Simulation Details

#### 1. Hy-Gō

CG molecular dynamics simulations of an uncapped heptamer of alanine residues (Ala_7_) with the Hy-Gō model were performed with the Gromacs 4.5.3 simulation suite^31^ in the constant NVT ensemble, while employing the stochastic dynamics algorithm with a friction coefficient *γ* = (2.0 𝒯^S^)^‒1^ and a time step of 1 × 10^‒3^ 𝒯^S^. For each model and for each temperature considered, 20 independent simulations were performed with starting conformations varying from full helix to full coil. Each simulation was performed for 100,000 𝒯^*S*^, recording the system every 0.5 𝒯^S^. The CG unit of time, 𝒯^S^, can be determined from the fundamental units of length, mass, and energy of the simulation model, but does not provide any meaningful description of the dynamical processes generated by the model. In this case, 𝒯^S^ = 1 ps.

#### 2. PLUM

CG simulations of Ala_7_ with the PLUM force field^14,25,32^ were run using the ESPResSo simulation package.^33^ For details of the force field, implementation, and simulation parameters, see Bereau and Deserno.^14^ For each temperature considered, a single canonical simulation was performed for 200,000 𝒯^P^ with at timestep of 0.01 𝒯^P^, recording the system every 0.5 𝒯^P^, where 𝒯^P^ ~ 0.1 ps. Temperature control was ensured by means of a Langevin thermostat with friction coefficient *γ* = (1.0 𝒯^P^)^‒1^.

#### 3. AA

As a preliminary reference to help guide our search of Hy-Gō parameter space, we employed AA simulations of Ala_7_, previously published by Stock and coworkers.^34^ In short, these simulations employed the GROMOS 45A3 force field^35^ along with the SPC water model^36^ to obtain an 800 ns trajectory, sampled every 1 ps, for Ala_7_ in the zwitterionic state at 300 K.

### C. Markov State Models

Markov state models (MSMs) are kinetic models that characterize the probability of transitioning between a finite set of microstates, chosen to represent the configuration space of the underlying system.^37–39^ The transition probabilities may be estimated directly from molecular dynamics trajectories via a Bayesian scheme to enforce relevant physical constraints. In the present work, MSMs are built from the Ala_7_ helix-coil trajectories generated from each simulation model. To determine the MSM microstate representation, Principle Component Analysis was performed on the configurational space characterized by the *ϕ*/*ψ* dihedral angles of each residue along the peptide backbone. A density clustering algorithm was then applied to the five “most significant” dimensions in order to determine the number and placement of microstates, following previous investigation of the AA trajectory.^40^ This procedure yields 32 states in all cases, corresponding to the enumeration of all possible helical/coil state combinations for each of the 5 peptide bonds. As described further in the Supporting Information section, the helical (h) and coil (c) states for individual peptide bonds roughly correspond to the *α* and *β* regions, respectively, of the corresponding Ramachandran plot. MSM construction and analysis was performed using the pyEmma package.^41^ (See Supporting Information for more details and MSM validation.) We also considered lower resolution MSMs, characterized by the number of helical residues in a conformation, *Q_h_*. These MSMs were constructed from the “full” MSM via a direct mapping of the transition probability matrix. Such a construction compromises the quantitative description of kinetics. However, we found that this reduced resolution provides a useful tool for assessing the differences between various kinetic descriptions of Ala_7_.

## III. RESULTS AND DISCUSSION

In this study, we investigate the relationship between structural and kinetic properties of helix-coil transition networks generated by microscopic simulation models. The equilibrium statistical mechanics of helix-coil transitions is well-characterized by a lD-Ising model, which represents the state of each residue as being either helical, h,or coil, c.^42^ These simple equilibrium models employ two parameters, *w* and *v*, according to the Lifson-Roig (LR) formulation,^43^ which are related to the free-energy of helix propagation and nucleation, respectively. These parameters may be determined directly from simulation data using a Bayesian approach,^44^ and describe the over-arching structural characteristics of the underlying ensemble (see Supporting Information for more details). Perhaps the most important quantity, the average fraction of helical segments, 〈*f*_h_〉, i.e., propensity of sequential triplets of h states along the peptide chain, may be measured directly from calorimetry experiments. The average fraction of helical residues, regardless of the particular sequence of states along the peptide chain, denoted 〈*N*_h_〉, provides a complimentary observable to 〈*f*_h_〉. Although residue‐ or sequence-specific LR parameters may be determined in order to more faithfully reproduce the helix-coil properties generated from a simulation of a particular peptide system, in this work we determine a single set of {*w*, *v*} for each simulation model. These simple models are incapable of describing certain features, e.g., end effects, of the underlying systems; however, we utilize the LR parameters only as a characterization tool and demonstrate that the sequence-independent parameters are sufficient to effectively distinguish between various characteristics of the underlying simulations.

The kinetics of helix-coil transitions are often interpreted in terms of a kinetic extension of the Ising model.^45,46^ These kinetic models typically assume a simple relationship between the LR parameters and the on/off rate of helix formation and have been widely successful in describing the emergent kinetic properties, e.g., folding/unfolding rates, observed in experiments.^46,47^ However, the kinetics of the helix-coil transition may demonstrate drastic divergences from this behavior, e.g., when misfolded intermediates complicate the network of transition pathways.^48^ Even in simpler cases, the precise impact of the model’s assumptions on the fine details of the resulting kinetic network is not well understood.

Here, we construct kinetic models directly from the simulation trajectories, i.e., Markov state models (MSMs), allowing a more complex relationship between the LR parameters and the kinetic properties of the system. A comparison of these MSMs with the more approximate kinetic models clarifies the role that the physical details of the simulation models play in restricting the kinetic properties that are attainable via molecular simulations.

As a model system, we consider an uncapped heptapeptide of alanine residues (Ala_7_), which has 5 peptide bonds. From the LR point of view, there are 32 states, determined by enumerating the various sequences of h’s and c’s. The focus of our study is a relatively simple, native-biased CG model. The hybrid Gō (Hy-Gō) model, is a flavored-Gō model,^16–19^ with native contact (nc) interactions (i.e., hydrogen-bonding-like interactions between *i*/*i* + 4 residues) and associated desolvation barriers (db) between *Ο*_*α*_ atoms, as well as generic hydrophobic (hp) attractions between all pairs of C*β* atoms. The relative strengths of these interactions are determined by the set of 3 model parameters: {*ϵ*_nc_, *ϵ*_db_, *ϵ*_hp_}. Fig. 1 presents a visualization of the model representation and also the corresponding interaction potentials. The model also employs physics-based interactions in the form of sterics (see bottom-right panel of Fig. 1) and torsional preferences along the backbone. We denote this model “hybrid” since it employs both traditional Gō-type interactions as well as detailed physics-based interactions determined from an AA model. Further details of the model and corresponding simulations are described in the Methods section.

**FIG. 1.**
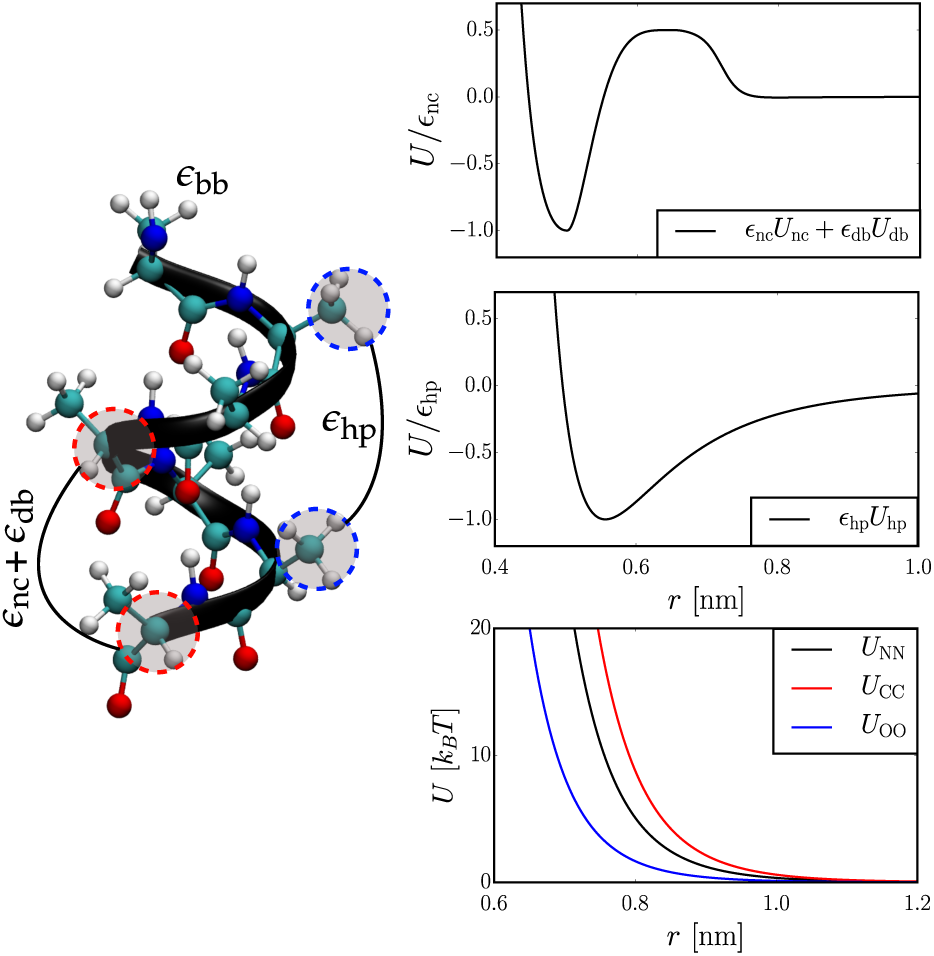
A visualization of the Hy-Gō model representation and interactions for Ala_7_. (Left) Illustration of a native contact between C_*α*_ atoms and a generic contact between C_*β*_ atoms, along with the corresponding parameters, (*ϵ*_nc_, *ϵ*_db_, *ϵ*_hp_}, associated with these interactions. (Right) The top two panels present the interaction potentials for the Gō-type interactions as a function of the model parameters. In the top panel, *ϵ*_db_ = 0.5*ϵ*_nc_. The bottom panel presents the Weeks-Chandler-Andersen-like potentials employed to model sterics along the peptide backbone.

To investigate potential relationships between structural and kinetic properties, we simulated Hy-Gō models with distinct combinations of *ϵ*_nc_, *ϵ*_db_, and *ϵ*_hp_ and then monitored various aspects of the resulting transition networks. As a preliminary reference to help guide our search of Hy-Gō parameter space, we employed an allatom (AA) simulation, previously studied by Stock and coworkers.^34,40,49^ A comparison between the Hy-Gō and AA trajectories additionally provides an assessment of the capability of the simple native-biased model to reproduce kinetic properties generated by a more detailed model. We stress that it is well-known that distinct AA force fields yield widely varying results, e.g., in terms of helical propensities, for short peptide systems.^44^ The AA simulation employed in this work serves as an ideal reference since it samples a largely disordered ensemble with a diverse collection of pathways from the coil to helix state and, thus, a complex hierarchy of relevant kinetic processes.

Fig. 2 presents a representative network description of the helix-coil trajectories examined in this study. The network is plotted along simple reaction coordinates that characterize the helicity of each conformation on the horizontal axis and the direction (c‐ or n-terminus) of folding on the vertical axis. The thickness of the arrows denote the relative probability flux passing between pairs of states for trajectories starting from the coil state and ending at the helix state. The “middle” of the network appears quite random, with more directed transitions towards the ends of the graph. Interestingly though, the residue dynamics remain highly coupled (see Fig. S4). The color of each state denotes its committor value—the probability of reaching the helix state before returning to the coil state. Clearly the landscape is largely tilted toward the unfolded region, as the committor values are very close to 0 for all but a few states.

**FIG. 2.**
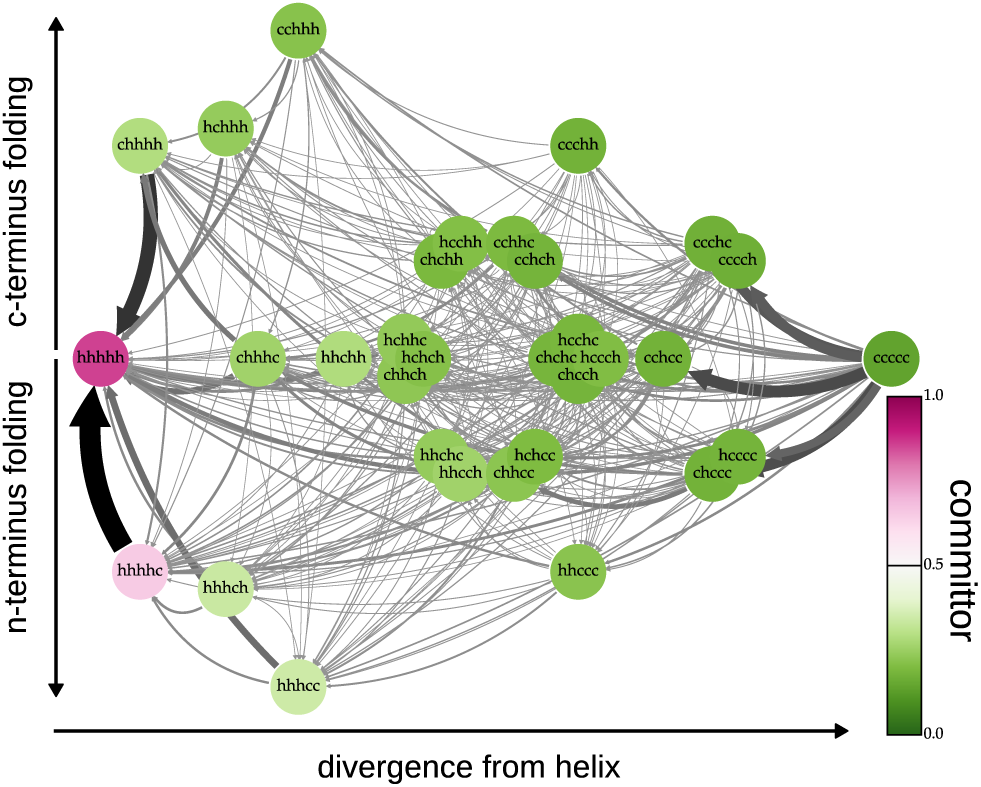
Network representation for the Ala_7_ helix-coil transition generated from an atomically-detailed simulation. The structural ensemble is representation by 32 microstates, corresponding to each possible sequence of the 5 peptide bonds attaining either a helical, h, or coil, c, state. The horizontal axis characterizes the difference between the sequence of each state and the native (full helix) state, while the vertical axis quantifies the extent to which the state corresponds to nor c-terminus folding. The thickness of the arrows represents the fractional probability flux passing through a pair of states for trajectories which begin in the coil state and end in the helix state. The color of each state denotes the value of the committor—the probability of reaching the helix state before returning to the coil state.

### A. Identifying structural-kinetic relationships at a single temperature using the Hy-Gō model

In order to assess the capability of the Hy-Gō model to reproduce structural and kinetic properties of the AA model, we considered various combinations of *ϵ*_nc_, *ϵ*_db_, and *ϵ*_hp_ and then adjusted the temperature to reproduce the average helicity of the AA model, 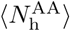. We found three classes of CG models: (*i*) those that reproduced 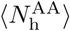 at some temperature *T*^*^, (*ii*) same as (*i*), but whose native state was the coil state, and (*iii*) those where the helix state was native but the model could not achieve such a low value of helicity at any temperature. For each model, we determined the two LR parameters, {*w*,*v*}, and constructed a 32-state Markov state model (MSM) directly from the simulation data.

Fig. 3 presents a “parameter landscape” for the Hy-Gō model, plotted as a ternary diagram with each axis characterizing the relative importance of the three Gō-type interactions: *ϵ*_*i*_ /*E*_tot_, where *E*_tot_ ≡ *ϵ*_nc_ + *ϵ*_db_ + *ϵ*_hp_. Because the relative impact of the backbone interactions depends on *E*_tot_, we discretize along *E*_tot_ and plot ternary diagrams for distinct values along this coordinate. Squares indicate models of type (*i*), circles of type (*ii*), and triangles of type (*iii*). The color of each model denotes the error with respect to some property of the AA model (cooler colors represent lower errors). The first row characterizes the root mean square error (rmse) with respect to the LR parameters of the AA model, while the second row quantifies the rmse with respect to the slowest two dynamical processes of the system (as characterized by the corresponding eigenvectors of the MSM at the *Q*_h_ level of resolution, see the Methods section for details) and also the ratio of timescales between these two processes. There is a clear correspondence between models that reproduce the structural and kinetic metrics. In other words, matching the overarching structural features of the ensemble (i.e., 〈N_h_〉 and 〈*f*_*h*_〉) is enough to also accurately reproduce the overarching hierarchy of kinetic processes of the underlying system at *T*^*^.

**FIG. 3.**
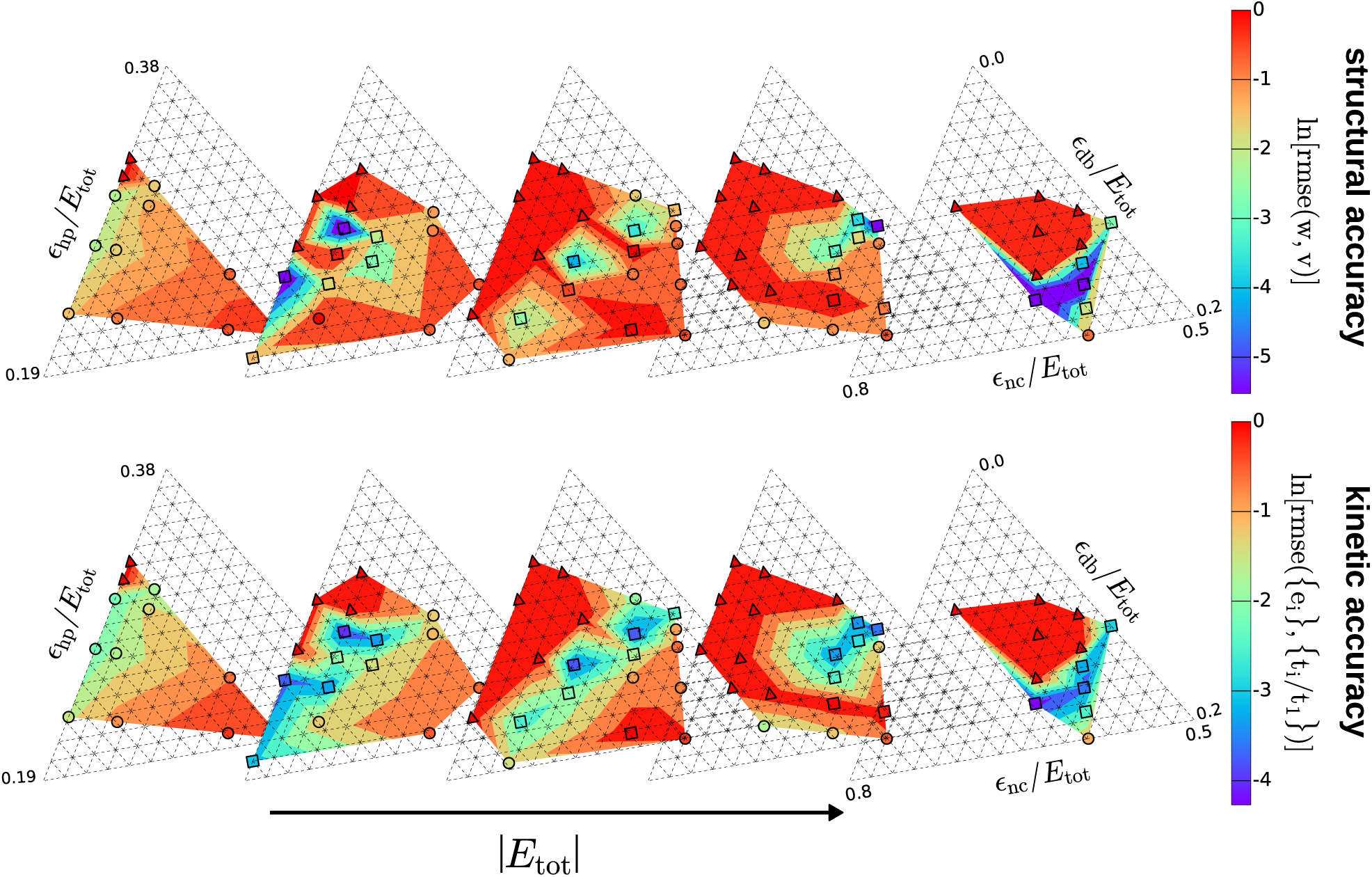
“Parameter landscape” for the Hy-Gō model. The axes of the ternary plots characterize the relative importance of each of the three Gō-type interactions: *ϵ*_i_ / *E*_tot_, where *E*_tot_ ≡ *ϵ*_nc_ + *ϵ*_db_ + *ϵ*_hp_. Because the relative impact of the backbone interactions depends on *E*_tot_, the plot is discretized along *E*_tot_. Squares indicate models that reproduced 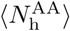 at some temperature *T*\*, circles indicate models similar to squares but whose native state is the coil state, and triangles indicate models with a helix native state but that could not achieve 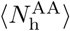 at any temperature. The color of each marker denotes the error with respect to some property of the AA model (cooler colors represent lower errors). Note that the filled color grid between points is included only to guide the eye. (Row 1) Root mean square error (rmse) with respect to the LR parameters of the AA model. (Row 2) rmse with respect to the slowest two dynamical processes of the system (as characterized by the corresponding eigenvectors of the MSM at the *Q*_h_ level of resolution) and also the ratio of timescales between these two processes. Because the errors of the models marked with triangles were considerably larger than the other models, their error values were reduced to the maximum of the other models for easier visual comparison.

Fig. 4 investigates the structural-kinetic relationships identified in Fig. 3 in greater detail. From the parameter search, 7 “AA-like” Hy-Gō models were identified that quantitatively reproduced 〈*N*_h_〉 and 〈*f*_h_〉 of the AA model. Fig. 4 presents various static and kinetic properties generated from the AA model (solid, black curves), the 7 AA-like Hy-Gō models (colored curves), and for two additional Hy-Gō models that also reproduce 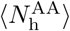 (while still sampling all 32 states), but do so at the largest and smallest simulated temperatures (dashed and dashed-dotted, black curves, respectively). The properties of the latter two models represent the range of attainable values of each observable, given the weak structural constraint of 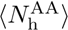 and also the implicit constraints enforced by the details of the model, but allowing for distinct LR parameters and, thus, –*f*_h_〉 values. Panel (*ai*) presents 〈*N*_h_〉 and 〈*f*_h_〉 (described above), as well as the average fraction of neighboring pairs of helical residues, 〈*N*_s_〉, and the average fraction of isolated helical residues, 〈*N*_l_〉. Panel (*aii*) presents the equilibrium distribution along the number of helical residues in the sequence, *Q*_h_. These panels demonstrate that the structural characteristics of the ensembles generated by the Hy-Gō models are largely fixed by fixing the average fraction of helical residues, 〈*N*_h_〉, and the average fraction of helical segments, 〈*f*_h_〉.

**FIG. 4.**
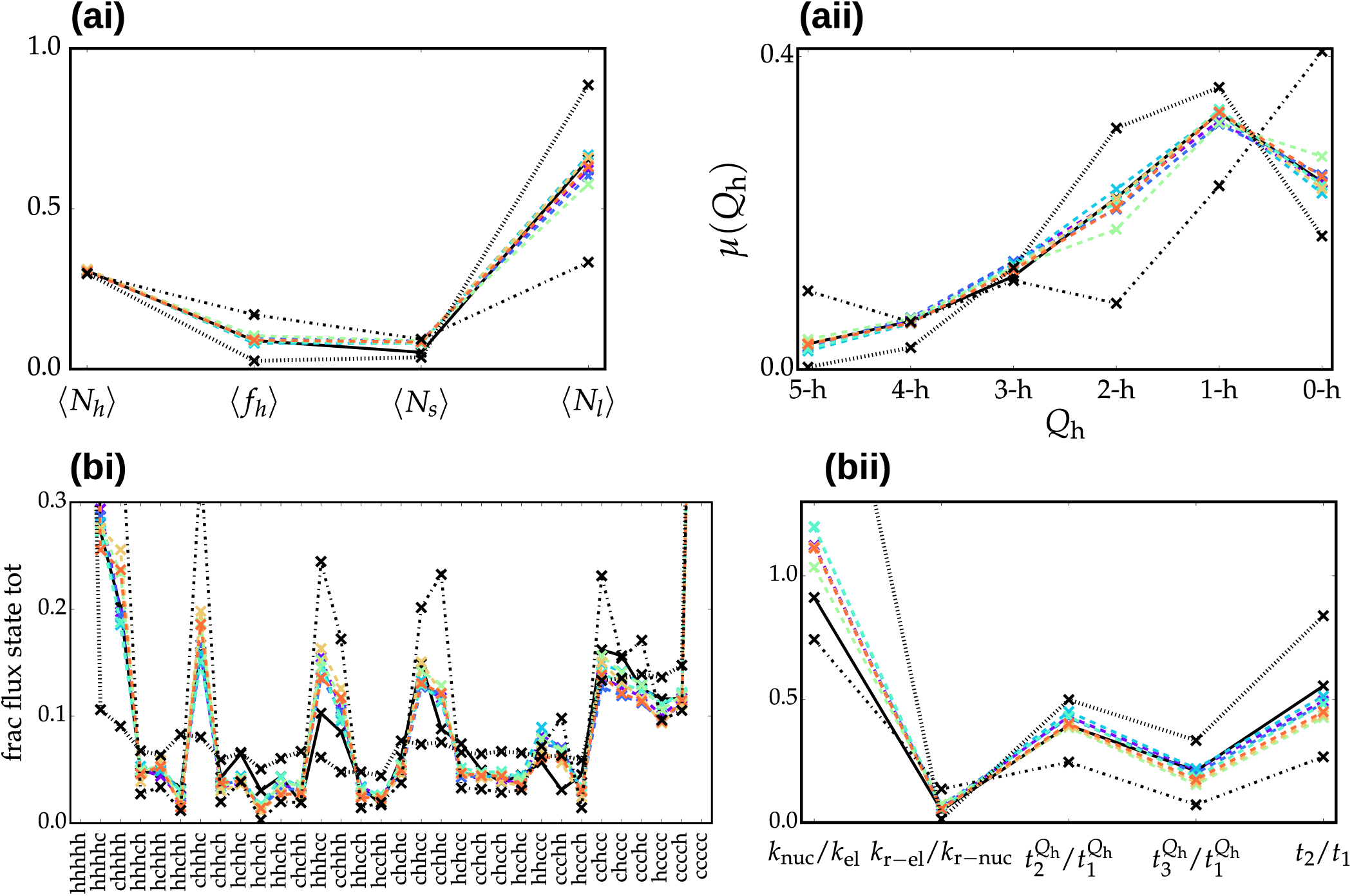
Static (row *a*) and kinetic (row *b*) properties for the Ala_7_ helix-coil transition generated by an all-atom model (AA; solid, black curves) and various Hy-Gō models. 7 AA-like Hy-Gōmodels (colored curves) were identified as optimally reproducing the properties of the AA model. The dashed and dashed-dotted black curves denote two Hy-Gō models that reproduce 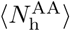 at the highest and lowest temperatures, respectively, of all models considered. (*ai*) The average fraction of helical residues, 〈*N*_h_〉, average fraction of helical segments, 〈*f*_h_〉, average fraction of sequential pairs of helical residues, 〈*N*_s_〉, and average fraction of isolated helical residues, 〈*N*_1_〉. (*aii*) The equilibrium distribution along the number of helical residues in the sequence, *Q*_h_. (*aiii*) The average fraction of helical segments per residue, 〈*h*(*i*)〉. (*bi*) The fractional flux passing through each state when considering pathways which begin in the coil state and end in the helix state. (*bii*) Ratios of important timescales. *k*_nuc_/*k*_el_ (*k*_r-nuc_/*k*_r-el_) is the ratio of (reverse) nucleation to (reverse) elongation rate. *t*_*i*_/*t*_1_ denotes the ratio of the *i*th slowest kinetic process to the slowest kinetic process according to the MSM. The *Q*_h_ superscript denotes timescales calculated at the resolution of the number of h residues in the sequence.

Panel (*bi*) presents the fractional flux passing through each state for trajectories that begin in the coil state and end in the helix state, while panel (*bii*) presents ratios of important timescales in the underlying simulations. It is important to note that, since the CG unit of time does not correspond to a physical time, there is always an arbitrary speed-up associated with dynamical properties generated by CG models. For this reason, we only consider ratios of timescales, to effectively account for this speed-up. *k*_nuc_ (*k*_r-nuc_) and *k*_el_ (*k*_r-el_) characterize the (reverse) rates of nucleation and elongation, respectively (see Supporting Information for details about the calculation of these rates). *t*_*i*_ denotes the timescale of the *i*^th^ slowest kinetic process according to the MSM. The Qh superscript denotes timescales calculated from a reduced MSM with microstates corresponding the number of h residues in the sequence. Panel (*b*) demonstrates that, quite remarkably, detailed kinetic features generated by the Hy-Gō models are also largely fixed by fixing 〈*N*_h_〉 and 〈*f*_h_〉. Interestingly, the approximate bounds on the timescale ratios, represented by the black dashed and dashed-dotted curves, demonstrate that there is a rather small amount of freedom in these quantities (other than for *k*_nuc_/*k*_el_). We suggest that this is due to the implicit constraints on the kinetic network, enforced by the underlying physics of the model.

Fig. 5 presents network representations of the AA model and a representative AA-like Hy-Gō model (top panel) as well as the low‐ and high-temperature Hy-Go models introduced above. The networks are presented in a different way than Fig. 1, with the horizontal axis corresponding to the value of the committor for each state and the vertical axis corresponding to the local graph entropy: *S*^loc^(*i*) = ∑_*j*_*T*_*jj*_ ln *T*_*jj*_, where *T*_*ij*_ is the transition probability from state *i* to state *j*. *S*^loc^ (*i*) quantifies the degree of randomness for pair transitions originating in state *i*. For visual clarity, we discretized the networks along these two coordinates and grouped states sharing the same grid into a single node. The relative size of each node denotes the number of underlying states. The color of each node corresponds to the average fractional flux passing through the node for trajectories starting from the coil state and ending in the helix state. The networks in Fig. 5 provide an illustration of the network topologies which generate the various properties presented in Fig. 4. The AA-like model appears to mimic the AA network in many ways, although the spread of states along each axis is not quite reproduced. On the other hand, the low‐ and high-temperature networks display drastically different topologies, albeit while retaining the average helicity of the AA model, 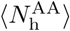. Thus, the average fraction of helical residues, 〈*N*_h_〉, is a rather weak structural constraint for this system.

**FIG. 5.**
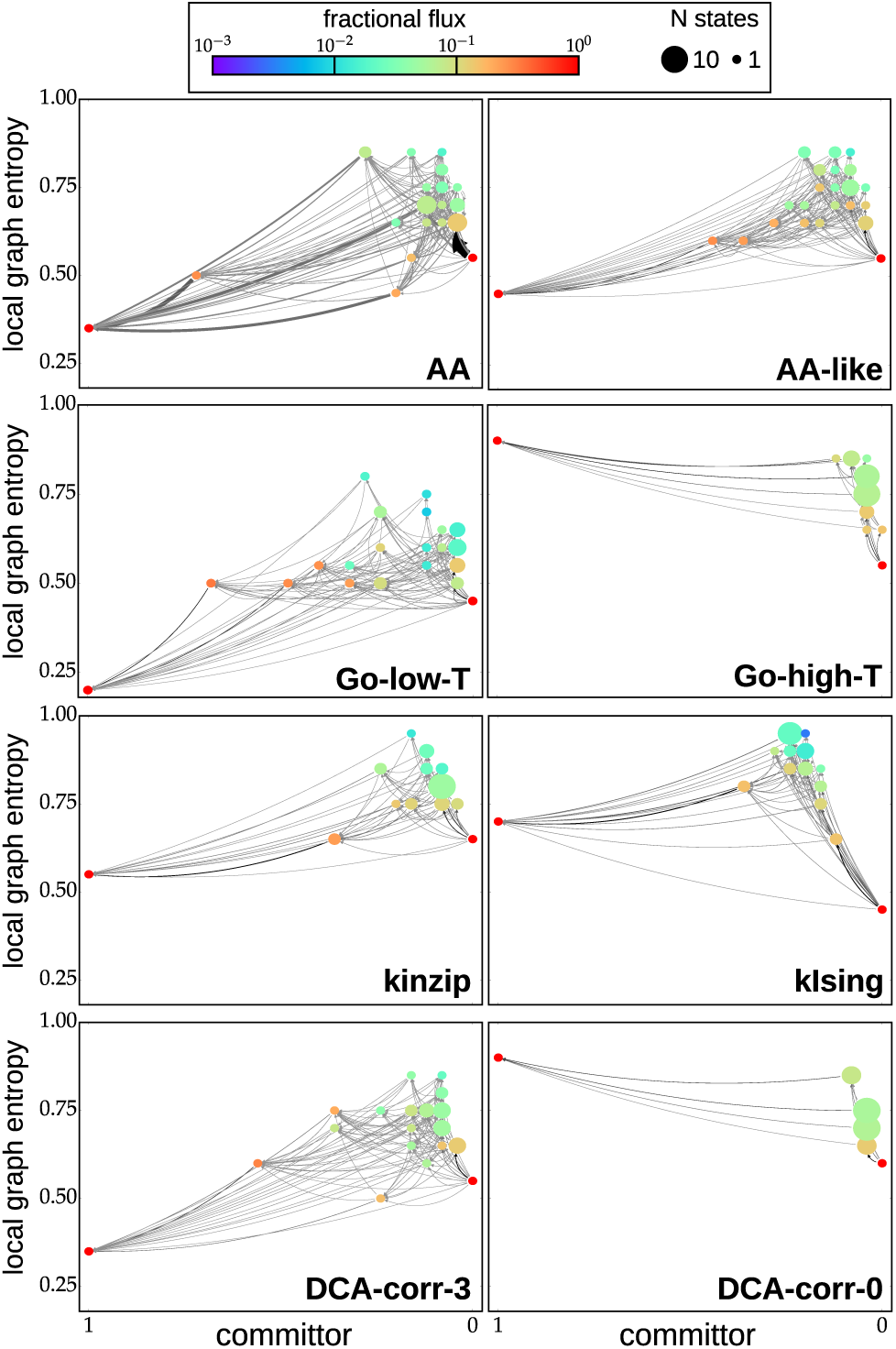
Network representations of the Ala_7_ helix-coil transition generated by an AA model, several CG models, and various approximations constructed from the AA simulation. (See main text for descriptions of the CG models and approximations). Each network is presented along the committor (i.e., the probability of reaching the helix state before returning to the coil state) on the horizontal axis and the local graph entropy, *S*^loc^, on the vertical axis. The local graph entropy is normalized by the maximum possible value (i.e., for a uniform distribution of transition probabilities). For visual clarity, we discretized the networks along these two coordinates and grouped states sharing the same grid into a single node. The relative size of each node denotes the number of underlying states. The color of each node corresponds to the average fractional flux passing through the node for trajectories from coil to helix.

### B. The impact of model physics on structural-kinetic relationships

In this section, we provide evidence that the structural-kinetic relationships identified in the previous section are due to the detailed physics built into the molecular simulation models. Approximate kinetic models built directly from the LR parameters provide a straightforward route to explicitly test the impact that the physical restrictions arising from the details of the molecular simulation models have on the resulting structural-kinetic relationships. We consider two models built directly from the LR parameters of the AA model, denoted kinzip and kIsing. The kinzip model corresponds to the kinetic zipper model,^46^ which assumes that the rate of transition from the c to h state for each residue is limited by the nucleation parameter, *v*. The kIsing model is based on the formulation of Schwarz,^45^ which uses the LR parameters to define the reaction rates for individual triplets of residues transitioning between conformational states. We also employed a different approach for constructing approximate models for the simulation kinetics, following the n-m approximation.^50^ In this procedure, we start with a “local” kinetic model determined from the AA simulation data, i.e., by ignoring the state of residues which lie beyond some number of peptide bonds away from a given residue, and then construct a full MSM, assuming that the residue dynamics beyond the chosen separation are decoupled. In this way, a systematic investigation into the coupling of residue dynamics along the peptide backbone can be performed. We refer to this procedure as the dynamic coupling analysis (DCA), and the corresponding models are denoted DCA-corr-x, where x is the chosen number of peptide bonds to be correlated in the model. See the Supporting Information section for details about the construction of each of these models.

Overall, the approximate kinetic models display varying structural and kinetic accuracy (Fig. S4), while reproducing 〈*N*_h_〉 and 〈*f*_h_〉. A full decoupling of residue dynamics (DCA-corr-0) introduces rather large errors into all other static and kinetic observables considered in Fig. 4. In general, decoupling of residue dynamics yields a spread of flux throughout the network and the systematic oversampling of intermediate states, as previously reported for bottom-up CG models for the helix-coil transition.^51^ The kIsing model displays deficiencies in particular static and kinetic properties, while the kinzip model demonstrates an accuracy for these observables that is on par with the AA-like Hy-Gō models. Fig. 5 presents network representations of the kinzip and kIsing models, and two approximate models constructed by systematic decoupling of the residue dynamics, assuming no correlations and *i*/*i*+3 correlations (DCA-corr-0 and DCA-corr-3,respectively). The kinzip and kIsing networks seem to reproduce the AA properties via a simplified network topology, while DCA-corr-3 retains a closer resemblance to the AA network. The DCA-corr-0 network displays drastically different topological features, as is apparent in the resulting properties of this model. Decoupling of residue dynamics leads to a decrease in committor values and an increase in local entropies, especially for the largely helical states.

These results provide evidence of the strong impact that the underlying physics of the Hy-Gō model has on the attainable static and kinetic properties generated by molecular simulations. Although the approximate kinetic models demonstrate varying network topologies, timescale ratios, and structural distributions while attaining the appropriate 〈*N*_h_〉 and 〈*f*_h_〉 values, Figs. 3-5 demonstrate that the properties of the Hy-Gō model are largely fixed for particular values of 〈*N*_h_〉 and 〈*f*_h_〉.

### C. Further validation of structural-kinetic relationships using a transferable CG model

We tested the robustness of the structural-kinetic relationships by considering the transferable CG PLUM model with three distinct parametrizations. The static and kinetic properties of the original PLUM model^14^ for Ala_7_ are presented in Fig. 6 as solid violet curves. The PLUM model strongly stabilizes helical structures for Ala_7_, and is incapable of achieving the low helical content of the AA model. The significant discrepancies between the AA (black, dashed-dotted curves) and PLUM model are not surprising since (*i*) neither the PLUM nor the AA model was parametrized to reproduce properties of small peptides and (*ii*) various AA models yield widely varying structural properties for peptide systems.^44^ Rather, in this study the AA and PLUM models represent two distinct reference ensembles for investigating the interplay between structure and kinetics. We extended the search in Hy-Gō parameter space to find a set of models which more closely reproduced the structural properties of the PLUM model. Fig. 6 demonstrates that by approximately matching 〈*N*_h_〉 and 〈*f*_h_〉, the “PLUM-like” Hy-Gō model nearly quantitatively reproduces both the equilibrium and kinetic properties of the PLUM model. The network topology of the PLUM-like Hy-Gō model also closely resembles that of the PLUM model (Fig. S8). While the PLUM model employs quite distinct and more complex energetic interactions compared with the Hy-Gō model, both models provide a near-atomistic representation of the peptide backbone. Thus, the results presented in this section further demonstrate the role of specific built-in physics in determining structural-kinetic relationships while also supporting the robustness of these relationships.

**FIG. 6.**
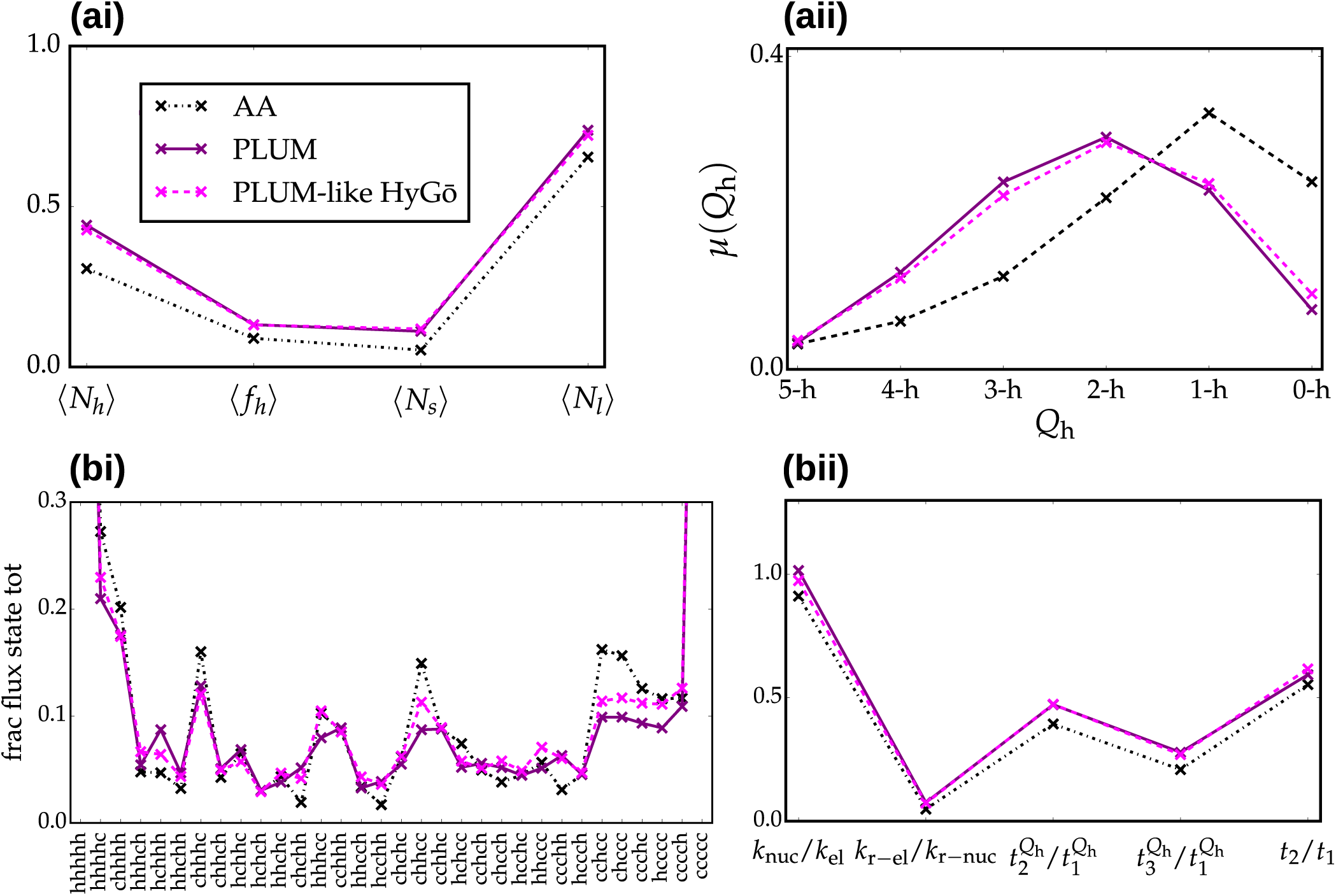
Static (row *a*) and dynamical (row *b*) properties for the Ala_7_ helix-coil transition generated by an all-atom model (AA; solid, black curves), the original PLUM model (PLUM; solid, violet curves), and a PLUM-like Hy-Gōmodel (PLUMlike; dashed, magenta curves). (*ai*) The average fraction of helical residues, 〈*N*_h_〉, average fraction of helical segments, 〈*f*_h_〉, average fraction of pairs of helical residues, 〈*N*_s_〉, and average fraction of isolated helical residues, 〈*N*_1_〉. (*aii*) The equilibrium distribution along the number of helical residues in the sequence, *Q*_h_. (*aiii*) The average fraction of helical segments per residue, 〈*h*(*i*)〉. (*bi*) The fractional flux passing through each state when considering pathways which begin in the coil state and end in the helix state. (*bii*) Ratios of important timescales. *k*_nuc_/*k*_el_ (*k*_r-nuc_/*k*_r-el_) is the ratio of (reverse) nucleation to (reverse) elongation rate. *t*_*i*_/*t*_1_ denotes the ratio of the *i*th slowest kinetic process to the slowest kinetic process according to the MSM. The *Q*_h_ superscript denotes timescales calculated at the resolution of the number of h residues in the sequence.

### D. Thermodynamics of Ala_7_ helix-coil transitions

To further validate the structural-kinetic relationships discussed above, we examined the temperature dependence of these relationships for each of the Hy-Gō and PLUM models. The temperature dependence of the helicity, in particular 〈*f*_h_〉(*T*), is a fundamental quantity for characterizing secondary structure formation in proteins and represents a simple measure for cooperativity of the helix to coil transition. We performed simulations of each model at various temperatures to determine 〈*f*_h_〉(*T*). The energy scales of the different models were aligned by shifting the temperature such that all models achieve 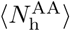 at *T*^*^. Although Fig. 4 demonstrates that the AA-like Hy-Gō models display very similar structural and kinetic properties at *T*_*_, Fig. S5 and Table S1 show that their cooperativities vary by more than 25% of the average value for these models. In other words, not surprisingly, reproducing simple structural properties at a single temperature is not enough to determine the thermodynamics of the model.

The robustness of the structural-kinetic relationships can be illustrated by considering the properties of all models, regardless of their particular structural features. Panel (*a*) of Fig. 7 presents the ratio of nucleation to elongation rates (for the forward and reverse processes) versus 〈*f*_h_〉 at *T*_*_ for the AA-like Hy-Gō models (purple, circle markers), other Hy-Gō models which reproduce 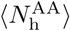 at some temperature *T*_*_ (blue, upward triangle markers), Hy-Gō models incapable of reproducing 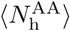 (green, sideways triangle markers), the PLUMlike Hy-Gō models (red, square markers), and the PLUM model (black, X markers). Panels (*bi*) and (*bii*) of Fig. 7 presents the slope of the linear fit of ln(*k*_nuc_/*k*_el_) and ln(*k*_r-nuc_/*k*_r-el_) as a function of inverse temperature, denoted Δ(*k*_nuc_/*k*_el_) and Δ(*k*_r-nuc_/*k*_r-el_), respectively, versus the slope of the linear fit of ln〈*f*_h_〉 as a function of inverse temperature, denoted Δ*f*_h_. There is a near one-to-one correspondence between each of the quantities presented in Fig. 7 (quantified by the Pearson correlation coefficient, R), although the forward rates show more complex behavior, with two clear regimes. These correlations demonstrate the robustness of the structural-kinetic relationships, which hold over largely varying structural ensembles and distinct model types.

**FIG. 7.**
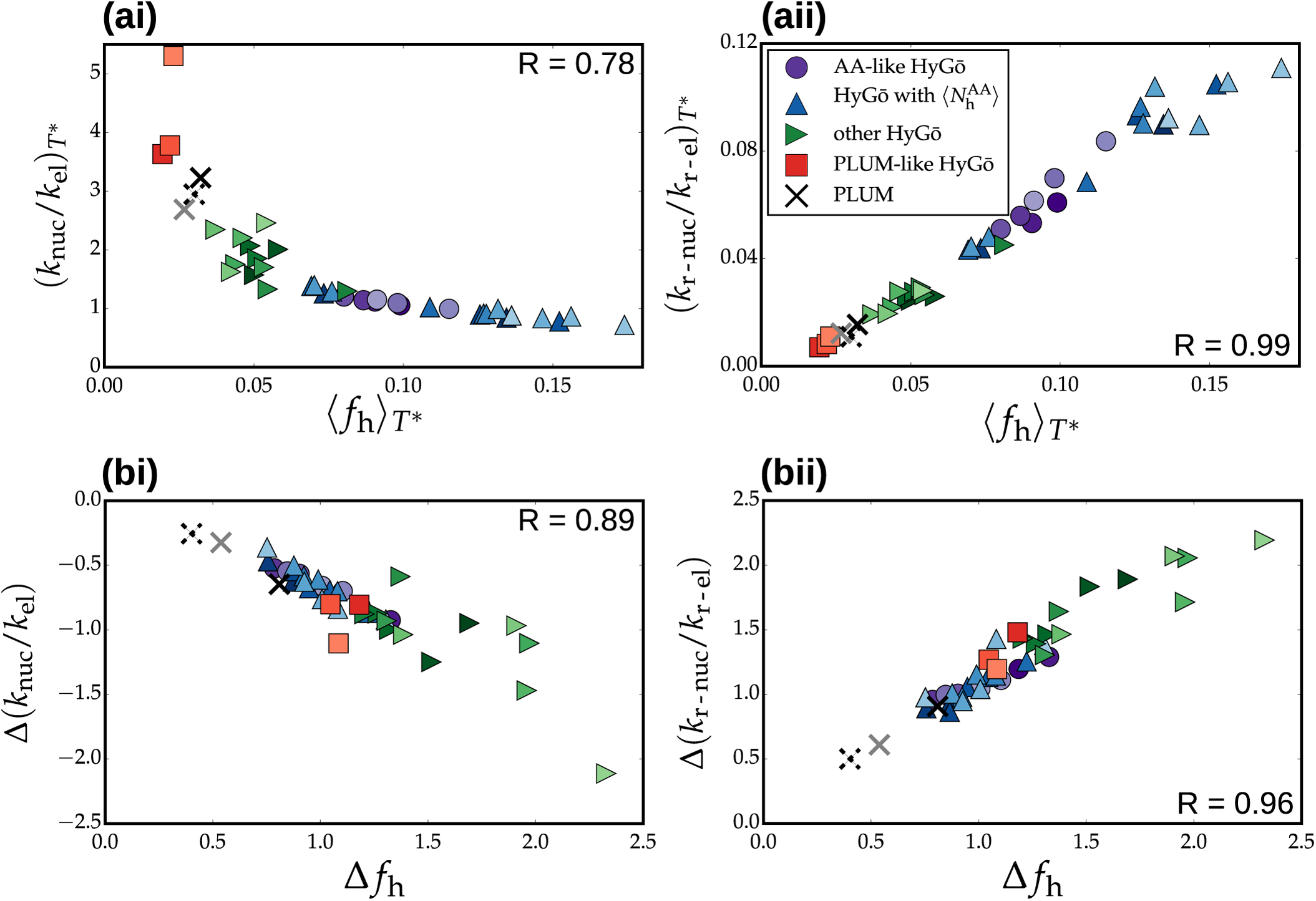
Relationships between static and kinetic properties. (*a*) Ratio of nucleation to elongation rate, *k*_nuc_/*k*_el_ (*ai*), and reverse rates, *k*_r-nuc_/*k*_r-el_ (*aii*) as a function of average fraction of helical segments, 〈*f*_h_〉, all determined at the reference temperature, *T*\*. (*b*) Slope of the linear fit of ln(*k*_nuc_/*k*_el_) (*bi*) and ln(*k*_r-nuc_/*k*_r-el_) (*bii*) as a function of inverse temperature, denoted Δ(*k*_nuc_/*k*_el_) and Δ(*k*_r-nuc_/*k*_r-el_), respectively, versus the slope of the linear fit of ln 〈*f*_h_〉 as a function of inverse temperature, denoted Δ*f*_h_. Colored circles, colored triangles, and open circles denote the AA-like Hy-Gōmodels, other Hy-Gō models, and the PLUM models, respectively. The PLUM models are denoted with X markers in the top row.

The particular relationships presented in Fig. 7 are unlikely to be generally valid for longer, heterogeneous sequences, where complex nucleation/elongation kinetics may arise.^48^ However, these results indicate that, when Arrhenius behavior holds, there may be simple relationships between the *T*-dependence of structural and kinetic properties. Moreover, these findings suggest that reproducing particular structural features over multiple temperatures may be an avenue for improving structural, thermodynamic and kinetic consistency for CG models.

### E. Relationships between thermodynamics and network topology

To conclude our investigation, we take a deeper look at the ensemble of pathways between helix and coil states, i.e., the topology of the kinetic networks generated by the various simulation models. The Markov state model characterization of the system kinetics provides a convenient and powerful framework for investigating the details of the hierarchy of kinetic processes sampled by particular helix-coil transitions. Indeed, graph-theoretic observables have been previously employed for understanding complex phenomena, e.g., kinetic frustration,^52^ arising within the dynamical processes of proteins. We applied discrete transition path theory^53^ to investigate the ensemble of pathways between the helix and coil states and identified simple relationships between the network topology and typical physical observables, e.g., the average fraction of helical segments, 〈*f*_h_〉. We found that 〈*f*_h_〉, at a particular temperature, is largely determined by the number of paths needed to account for half of the fractional flux of probability for trajectories traveling from the coil to the helix (Fig. S12). This somewhat surprising relation is likely due to the relatively strong constraints on the network topology enforced implicitly through the underlying physics of the models.

As the temperature of each system is decreased, the committor value of each state tends to increase while the local entropy tends to decrease (Fig. S11). Because of these generic changes in the network as a function of temperature, there may exist a feature of the network topology, at a particular reference temperature *T*^ref^, which determines the thermodynamics, i.e., cooperativity, of the corresponding transition. In the following, *T*^ref^ corresponds to the simulated temperature for each model that is closest to *T*_*_ (see Fig. S5). This slight inconsistency between the reference temperatures presumably introduces small errors into the following analysis. We consider the path entropy, i.e., Shannon entropy of the probability distribution of pathways, which characterizes the degree of randomness of paths traveling from a starting state, *s*, to an ending state, *d*. Similarly, the conditional path entropy,^54^ *H*_*sd*_|_*u*_, characterizes the average degree of randomness for paths from *s* to *d* passing through a particular intermediate state, *u*.

Naively, one may hypothesize that networks with more directed flux from coil to helix at *T*^ref^ would display a faster change of population into the helical state as the temperature is reduced. In other words, in order to maximize the cooperativity, the conditional path entropy, averaged over all intermediate states, should be minimized. To investigate the relationship between directed flux in the network and cooperativity of the underlying transition, we performed a correlation analysis by constructing a linear combination of input features, {f}, to characterize the target observable, *O*: *O* ≈ *F*[f] = ∑_*i*_c_*i*_f_*i*_, where the coefficients, {c_*i*_}, are determined as a best fit over all models. In this case, *O* corresponds to the slope of ln〈*f*_h_〉(1/*T*), denoted simply Δ*f*_h_. The set of 60 conditional path entropies for each intermediate state between coil and helix and for both the folding and unfolding directions were initially considered as input features. For numerical convenience, we weighted each of the features by the fractional flux passing through each state (as introduced above).

By systematically disregarding features which played a minimal role in the resulting correlation, we determined a reduced set of 16 features necessary to retain a reasonably accurate correlation (Fig. 8). Interestingly, the most important features corresponded to the conditional path entropies for a set of 8 states with intermediate helical content: {cchhh,hcchh,chhcc,cchhc,hchcc,chcch,hcchc,hhccc}.

**FIG. 8.**
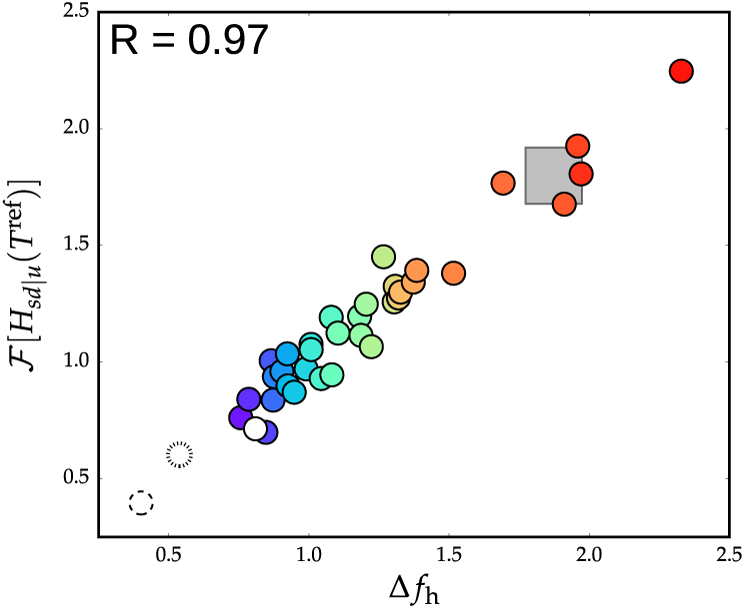
Relationships between thermodynamic and network properties. Correlation analysis using a reduced set of 16 features corresponding to the conditional path entropy, *H*_*sd*_|_*u*_, for each state in either the folding or unfolding direction for 8 particular states of intermediate helical content. Hy-Gō models are denoted by the colored circles, while the PLUM models are denoted by the empty circles. The gray box represents the predicted cooperativity of the AA model, based on the topological features of the network at *T*_*_.

This correlation analysis revealed that, unlike our naive assumption, it is not a simple minimization of the average conditional path entropies which determines a maximally cooperative model. Rather, there is a balance between achieving small conditional path entropies (i.e., directed flux) for some intermediate states and higher conditional path entropies (i.e., nondirected flux) for others (see Figs. S13 and S14 for more details). While the structural-kinetic relationships identified in previous sections probe the relationship between the physics of the simulation model and the shape of the free-energy landscape at a particular thermodynamic state point, the relationship between network topology at a single temperature and the cooperativity of the model suggests restricted changes in the free-energy landscape with respect to thermodynamic perturbations.

## IV. CONCLUSIONS

In this study, we identified robust relationships between the structural properties generated by a molecular simulation model and features of the corresponding hierarchy of kinetic processes. By considering simple kinetic models, we explicitly demonstrated that this relationship is dependent upon the implicit constraints enforced by the physics of the simulation model. In other words, the physics built into a simulation model shapes the free-energy landscape and constrains the allowable kinetic properties for any particular parametrization. Our previous work demonstrated that an accurate representation of local free-energy barriers is necessary for a faithful description of the hierarchy of kinetic processes.^4,6^ The results of the present work demonstrate, more generally, that when the essential physics is accounted for, i.e., when the structural ensemble is reproduced with sufficient accuracy, simple relationships between structure and kinetics may emerge.

The structural-kinetic relationships found in this work were identified through a systematic parameter scan which targeted the features of distinct reference models. We considered the helix-coil transition of a small peptide as a model system and employed a large variety of Hy-Gō models with varying interaction parameters and also three different parametrizations of the transferable CG PLUM model. The Hy-Gō models not only produced consistent kinetics by matching certain structural features of a reference model, but also demonstrated rather stringent topological constraints on the network, due to the underlying physics of the model. These constraints revealed an extension of the structural-kinetic relationships to thermodynamic quantities—the network topology at a single reference temperature determines the co-operativity of the resulting transition. While validation of particular structural-kinetic and network topology-thermodynamic relationships for, e.g., distinct peptide systems and tertiary structure formation remains for future work, the results presented here demonstrate a general approach for building kinetically-consistent CG models through the exploitation of structural-kinetic relationships. Therefore, our work motivates further investigation into structural-kinetic relationships for CG models and suggests that the matching of particular structural properties over multiple temperatures (or other thermodynamic state points) may provide a general scheme for simultaneous inclusion of structural, thermodynamic, and kinetic consistency into CG models.

## SUPPLEMENTARY MATERIAL

An online supplement to this article with further methodological details as well as additional results can be found at http://biorxiv.org.

## DATA

An onlinedatabaseconsisting ofvarious analysisscripts andinput files for the coarse-grainedsimulationscan be foundat https://github.com/JFRudzinski/Scripts_and_models_for_Structural-kinetic-thermodynamic_relationships.git

## ACKNOWLEDGEMENTS

We thank Fabian Knoch and Raffaello Potestio for critical reading of the manuscript. We are also very grateful to Gerhard Stock for the use of the Ala_7_ AA trajectory. J.F.R. thanks Florian Sittel for many fruitful discussions and assistance with the density clustering software. J.F.R. also thanks Amedeo Caflisch and Andreas Vitalis for useful discussions concerning the Lifson-Roig models. J.F.R. is grateful to the organizers of the Hunfeld workshop for supplying a stimulating atmosphere for scientific discussions which made a positive impact to this work. This work was funded through a postdoctoral fellowship from the Alexander von Humboldt foundation (J.F.R.) and an Emmy Noether fellowship of the Deutsche Forschungsgemeinschaft (T.B.).

